# The Relationship between Birth Timing, Circuit Wiring, and Physiological Response Properties of Cerebellar Granule Cells

**DOI:** 10.1101/2021.02.15.431339

**Authors:** S. Andrew Shuster, Mark J. Wagner, Nathan Pan-Doh, Jing Ren, Sophie M. Grutzner, Kevin T. Beier, Tony Hyun Kim, Mark J. Schnitzer, Liqun Luo

## Abstract

Cerebellar granule cells (GrCs) are usually regarded as a uniform cell type that collectively expands the coding space of the cerebellum by integrating diverse combinations of mossy fiber inputs. Accordingly, stable molecularly or physiologically defined GrC subtypes within a single cerebellar region have not been reported. The only known cellular properties that distinguishes otherwise homogeneous GrCs is the correspondence between GrC birthtime and the depth of the molecular layer to which their axons (parallel fibers) project. To determine the role birth timing plays in GrC wiring and function, we developed genetic strategies to access early- and late-born GrCs. We initiated retrograde monosynaptic rabies virus tracing from control, early-born, and late-born GrCs, revealing the different patterns of mossy fiber input to GrCs in vermis lobule 6 and simplex, as well as to early- and late-born GrCs of vermis lobule 6: sensory and motor nuclei provide more input to early-born GrCs, while basal pontine and cerebellar nuclei provide more input to late-born GrCs. *In vivo* multi-depth 2-photon Ca^2+^ imaging of parallel fibers of early- and late-born GrCs revealed representations of diverse task variables and stimuli by both populations, with differences in the proportions of parallel fibers encoding movement, reward anticipation, and reward consumption. Our results suggest neither organized parallel processing nor completely random organization of mossy fiber→GrC circuitry, but instead a moderate influence of birth timing on GrC wiring and encoding. Our imaging data also suggest that GrCs can represent general aversiveness, in addition to recently described reward representations.

**Significance Statement:** Cerebellar granule cells (GrCs) comprise the majority of all neurons in the mammalian brain and are usually regarded as a uniform cell type. However, the birth timing of an individual GrC dictates where its axon projects. Using viral-genetic techniques, we find that early- and late-born GrCs receive different proportions of inputs from the same set of input regions. Using *in vivo* multi-depth 2-photon Ca^2+^ imaging of axons of early- and late-born GrCs, we found that both populations represent diverse task variables and stimuli, with differences in the proportions of axons in encoding of a subset of movement and reward parameters. These results indicate that birth timing contributes to the input selection and physiological response properties of GrCs.

## Introduction

Cerebellar granule cells (GrCs) comprise the majority of neurons in the mammalian brain (1, 2). Each GrC receives only four excitatory inputs from mossy fibers, which originate in a variety of brainstem nuclei and the spinal cord, and the vast number of GrCs permits diverse combinations of mossy fiber inputs. Classical theories of cerebellar function have therefore proposed that GrCs function to integrate diverse, multimodal mossy fiber inputs and thus collectively expand coding space in the cerebellum (3–5). Until recently, studies have focused on the role of GrCs in implementing sparse coding of sensorimotor variables and stimuli (6–9). However, recent physiological studies of GrCs in awake, behaving animals highlight GrC encoding of cognitive signals in addition to sensorimotor signals (10–13). GrCs have also been recently shown to encode denser representations than expected by classical theory (10–12, 14–18), including a lack of dimensionality expansion under certain conditions (18).

Despite the vast number of GrCs, stable molecularly or physiologically defined GrC subtypes within a single cerebellar region or lobule have not been described (19–22). The only known axis along which spatially intermingled GrCs can be distinguished from each other is the depth of the molecular layer to which their parallel fiber axons (PFs) project, which is dictated by GrC lineage and birth timing (23, 24). Birth timing predicts the wiring and functional properties of diverse neuron types in many neural systems (25), including the neocortex (26, 27), other forebrain regions (28, 29), olfactory bulb (30, 31), and ventral spinal cord (32, 33). Furthermore, classic studies utilizing γ-irradiation at different times during rat postnatal development to ablate different cerebellar GrC and interneuron populations suggested that GrCs born at different times could contribute differentially to motor vs. action coordination (34). These observations also led to an as-of-yet untested hypothesis that mossy fibers arriving at different times during development could connect with different GrC populations. Could GrC birth timing be an organizing principle for information processing in the cerebellum?

Recent evidence and modeling point to the possibility of spatial clusters of co-activated PFs (15, 35), suggesting that GrCs born around the same time may disproportionally receive co-active mossy fiber inputs. However, another study using different methods and stimuli did not find differences in the physiological responses of early- and late-born GrCs to various sensorimotor stimuli (36). Here, we aimed to address the role of birth timing in GrC wiring and function. To do so, we developed strategies to gain genetic access to early- and late-born GrCs, as well as control GrCs not restricted by birth timing. We report the first monosynaptic input tracing to GrCs, finding differential mossy fiber inputs to GrCs in vermis lobule 6 and simplex, as well as different patterns of input to early- and late-born GrCs in vermis lobule 6. Finally, we performed *in vivo* multi-depth 2-photon Ca^2+^ imaging of PFs of early- and late-born GrCs during an operant task and presentation of a panel of sensory, appetitive, and aversive stimuli. We found differences in the proportions of early- and late-born GrCs encoding of a subset of movement and reward parameters. Together, these results reveal the contribution of GrC birth timing to their input wiring and diverse encoding properties.

## Results

### Genetic Strategies for Accessing Birth Timing-defined Cerebellar Granule Cells

Cerebellar cortical circuit assembly occurs primarily during the first three postnatal weeks in the mouse, reaching full maturity shortly thereafter (37). At birth, GrC progenitors occupy the most superficial external granular layer (EGL), where they proliferate. Mossy fibers are morphologically recognizable after postnatal day 5 (P5), shortly after GrC progenitors have begun to exit mitosis (38). Newborn GrCs extend their axons as PFs into the developing molecular layer between the EGL and Purkinje cell bodies. Newborn GrC somata descend past the Purkinje cell layer (PCL) into the internal granular layer (IGL), giving rise to the granule cell layer in adults (39). Later-born GrCs stack their PFs superficially to those of earlier-born GrCs (24). As GrC neurogenesis proceeds, the EGL is gradually replaced by the molecular layer until P21, when GrC neurogenesis is complete (Fig. 1A).

**Fig. 1.**
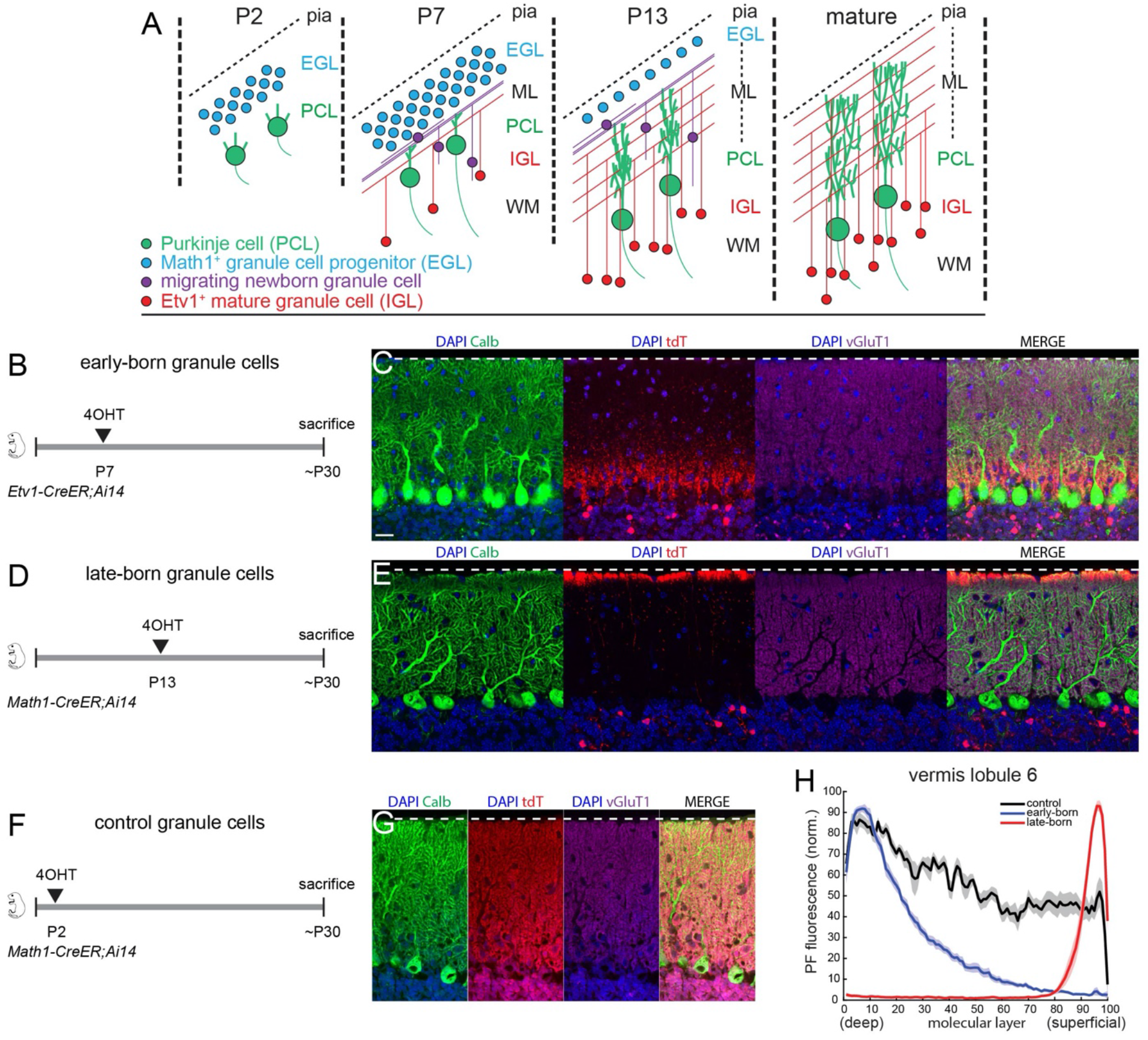
Genetic strategies for accessing birth timing-defined cerebellar granule cells (GrCs). (A) Schematic summary of postnatal GrC development. Math1 is expressed exclusively in GrC progenitors, and Etv1 in postmitotic GrCs. EGL, external granule cell (germinal) layer; pia, pia mater; ML, molecular layer; PCL, Purkinje cell layer; IGL, internal granule cell layer; WM, white matter layer. (B–G) Genetic strategies and corresponding images showing parallel fiber (PF) innervation of the cerebellar molecular layer for access to early-born (B, C), late-born (D, E), and control (F, G) GrCs. Images were taken from vermis lobule 6. Blue, DAPI; green, calbindin, a marker of Purkinje cells; red, tdTomato from the Cre reporter *Ai14*; purple, vGluT1, a marker of PF synapses. Dotted white lines indicate pia surface. Scale bar, 20 μm. (H) Quantification of PF-derived fluorescence across the depths of the molecular layer in vermis lobule 6. N = 12 sections from 2 mice each.

To gain access to early- and late-born GrCs for anatomical profiling and physiological recordings, we developed genetic strategies for accessing birth timing-defined GrCs. We scoured the Allen Brain Atlas transgenic characterization database for CreER lines with expression specific to GrC progenitors and mature GrCs, but not mossy fiber origin sites in the brainstem (40). We identified two lines designed to mimic the expression pattern of the transcription factors *Math1* (41), which is expressed selectively in GrC progenitors in the cerebellum (42), and *Etv1* (43), which is expressed selectively in postmitotic cerebellar GrCs. To test our genetic strategies, we crossed *Etv1-CreER* and *Math1-CreER* mice to tdTomato Cre reporter mice (44). We injected 4-hydroxytamoxifen (4OHT), the active metabolite of tamoxifen that drives CreER nuclear translocation, into *Etv1-CreER;Ai14* mice at P7 and *Math1-CreER;Ai14* mice at P13 to label early- and late-born, respectively, GrCs. As predicted, these strategies resulted in PF fluorescence selectively in the deep and superficial portions of the molecular layer, respectively (Fig. 1B–E), allowing selective genetic access to early- and late-born GrCs. Furthermore, injecting 4OHT into *Math1-CreER;Ai14* mice at P2 resulted in labeling of GrCs with PFs projecting to all depths of the molecular layer, indicating genetic access to GrCs not defined by birth timing as a control for comparison with birth timing-dependent results (Fig. 1F and G). Quantitative analysis of PF labeling showed that early- and late-born labeling strategies consistently allowed genetic access to specific and mutually exclusive populations of granule cells, but that the early-born strategy had a longer “tail” and thus more overlap with cells accessed via the control strategy (Fig. 1H).

### Differential Mossy Fiber Inputs to Granule Cells in Vermis Lobule 6 and Simplex

To our knowledge, mossy fiber inputs to GrCs from different precerebellar nuclei have not been quantitatively compared, largely due to longstanding technical limitations in performing input tracing from densely packed GrCs. Thus, prior to comparing inputs to early- and late-born GrCs, we first profiled the presynaptic inputs of mossy fibers to birth timing-unbiased GrC populations. We applied monosynaptic retrograde rabies virus (RV) tracing (Wickersham et al., 2007; Callaway and Luo, 2015) of mossy fiber input to GrCs of vermis lobule 6 and simplex. To do so, we crossed *Math1-CreER* mice to *R26^CAG-LSL-HTB^* (*R26^HTB^* hereafter) mice (45), which express rabies G protein (B19SAD-RG) and the EnvA receptor TVA in a Cre-dependent manner. We injected *Math1-CreER*;*R26^HTB^* mice with 4OHT at P2 and then injected G-deleted, GFP-expressing, EnvA-pseudotyped RV into vermis lobule 6 or simplex (bilateral) around P30 (P28–33; Fig. 2A). We sacrificed animals 5 days later and quantified GFP-labeled cells (Fig. 2B). This analysis revealed inputs from cells in many regions known to originate mossy fibers (46, 47), including the pons, various sensory and motor brainstem nuclei, and the cerebellar nuclei (Fig. 2C–M). In the absence of Cre, there were no retrogradely-labeled cells in the brainstem (n = 8 mice); by contrast, the brainstems of experimental animals had thousands of input cells labeled (vermis: 2285 ± 262 cells; simplex: 2746 ± 371 cells), validating the specificity our transgenic-viral tracing methods (Fig. S2A).

**Fig. 2.**
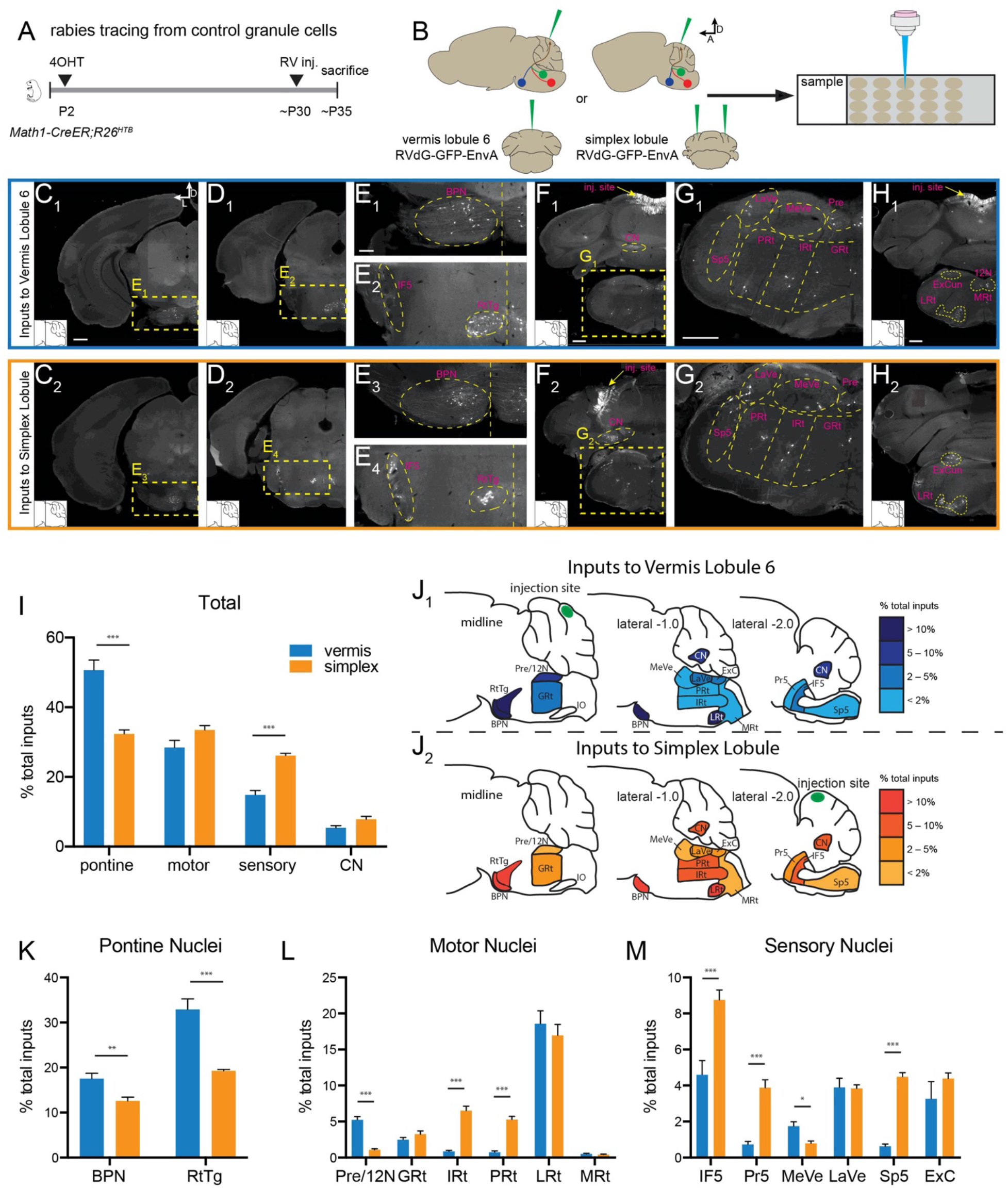
Rabies tracing of mossy fiber inputs to GrCs of vermis lobule 6 and simplex. (A, B) Time course (A) and schematic (B) of retrograde monosynaptic rabies tracing. RVdG-GFP-EnvA, rabies virus deleted for glycoprotein, expressing GFP, and pseudotyped with EnvA. D, dorsal; A, anterior. Top, sagittal view; bottom, coronal view. (C–H) Example GFP-labeled input cells to vermis lobule 6 (C_1_–H_1_) and simplex (C_2_–H_2_) GrCs. D, dorsal; L, lateral. Scale bars, 500 μm for C, D, F, G, H; 200 μm for E. Insets at the bottom left show section planes in sagittal atlas. (I) Total distribution of presynaptic inputs from super-categorized brainstem nuclei to vermis lobule 6 and simplex GrCs. (J) Schematic summary of distribution of presynaptic inputs from brainstem nuclei to vermis lobule 6 and simplex GrCs. (K–M) Distribution of inputs from brainstem pontine (K), motor (L), and sensory (M) nuclei to vermis lobule 6 and simplex GrCs. N = 8 (vermis), 8 (simplex). Error bars, SEM. \**p* < 0.05, \*\**p* < 0.01, \*\*\**p* < 0.001 (multiple unpaired t tests with Bonferroni correction). BPN, basal pontine nucleus; CN, cerebellar nuclei; ExC, external cuneate nucleus GRt, gigantocellular reticular nucleus; IF5, interfascicular trigeminal nucleus; IRt, intermediate reticular nucleus; LaVe, lateral vestibular nuclei; LRt, lateral reticular nucleus; MeVe, medial vestibular nuclei; MRt, medial reticular nucleus; Pr5, principle trigeminal sensory nucleus; Pre/12N, prepositus/hypoglossal nucleus; PRt, parvicellular reticular nucleus; RtTg, reticulotegmental nucleus of the pons; Sp5, spinal trigeminal nucleus.

We grouped inputs into four super-categories: pontine, motor, sensory, and cerebellar nuclei (CN; see Methods). Pontine input comprises input from the basal pons (BPN) and reticulotegmental nucleus of the pons (RtTg), both of which relay cortical information. Motor nuclei include the prepositus/hypoglossal nuclei (Pre/12N) and the gigantocellular (GRt), intermediate (IRt), parvicellular (PRt), lateral (LRt), and medullary (MRt) nuclei of the brainstem reticular system. Sensory nuclei include the interfascicular (IF5), primary sensory (Pr5), and spinal (Sp5) trigeminal nuclei, the medial (MeVe) and lateral (LaVe) vestibular nuclei, and the external cuneate nucleus (ExC). Among inputs to both vermis lobule 6 and simplex, very few spinal cord and inferior olive (IO) cells (≪1%, data not shown) were labeled, and so were not included in subsequent quantifications.

At the super-category level, quantitative analysis revealed that vermis lobule 6 received proportionally more pontine input than did simplex, while simplex received proportionally more input in aggregate from sensory brainstem nuclei than did lobule 6 (Fig. 2I). Quantitative analysis of individual nuclei revealed differential contributions to lobule 6 and simplex (Fig. 2J–M). For example, within the motor nuclei, lobule 6 received much more input from Pre/12N than did simplex, while simplex received more input from IRt and PRt than did lobule 6 (Fig. 2L). Among inputs from sensory nuclei, simplex received more input from trigeminal nuclei (IF5, Pr5, Sp5) than did lobule 6, while lobule 6 received more input from MeVe than did simplex (Fig. 2M). Finally, lobule 6 received a higher proportion of total inputs from pontine nuclei (Fig. 2K), in contradiction to classical conceptions of input to vermis coming mostly from the body rather than the cortex (48), but in line with recent multisynaptic tracing in rodents revealing substantial cerebral cortical input to lobule 6 (18, 49) and dorsal vermis generally (50). These results also suggest that monosynaptic retrograde rabies virus tracing is sensitive enough to reveal quantitative differences in mossy fiber distributions.

### Differential Mossy Fiber Inputs to Early- and Late-born Lobule 6 Granule Cells

We next performed monosynaptic retrograde RV tracing to reveal the distributions of mossy fiber input to early- and late-born GrCs. We crossed *Math1-CreER* and *Etv1-CreER* mice to *R26^HTB^* mice, injected *Etv1-CreER*;*R26^HTB^* mice and *Math1-CreER*;*R26^HTB^* with 4OHT at P7 and P13, respectively, to gain access to early- and late-born GrCs, followed by injecting vermis lobule 6 with RVdG-GFP-EnvA. We sacrificed mice 5 days later and analyzed rabies labeling results (Fig. 3A). We detected fewer input cells from early- and late-born samples than control samples (early-born: 476 ± 54 cells; late-born: 440 ± 56 cells), in accordance with the proportion of granule cells genetically accessed by each strategy (Fig. S3D).

**Fig. 3.**
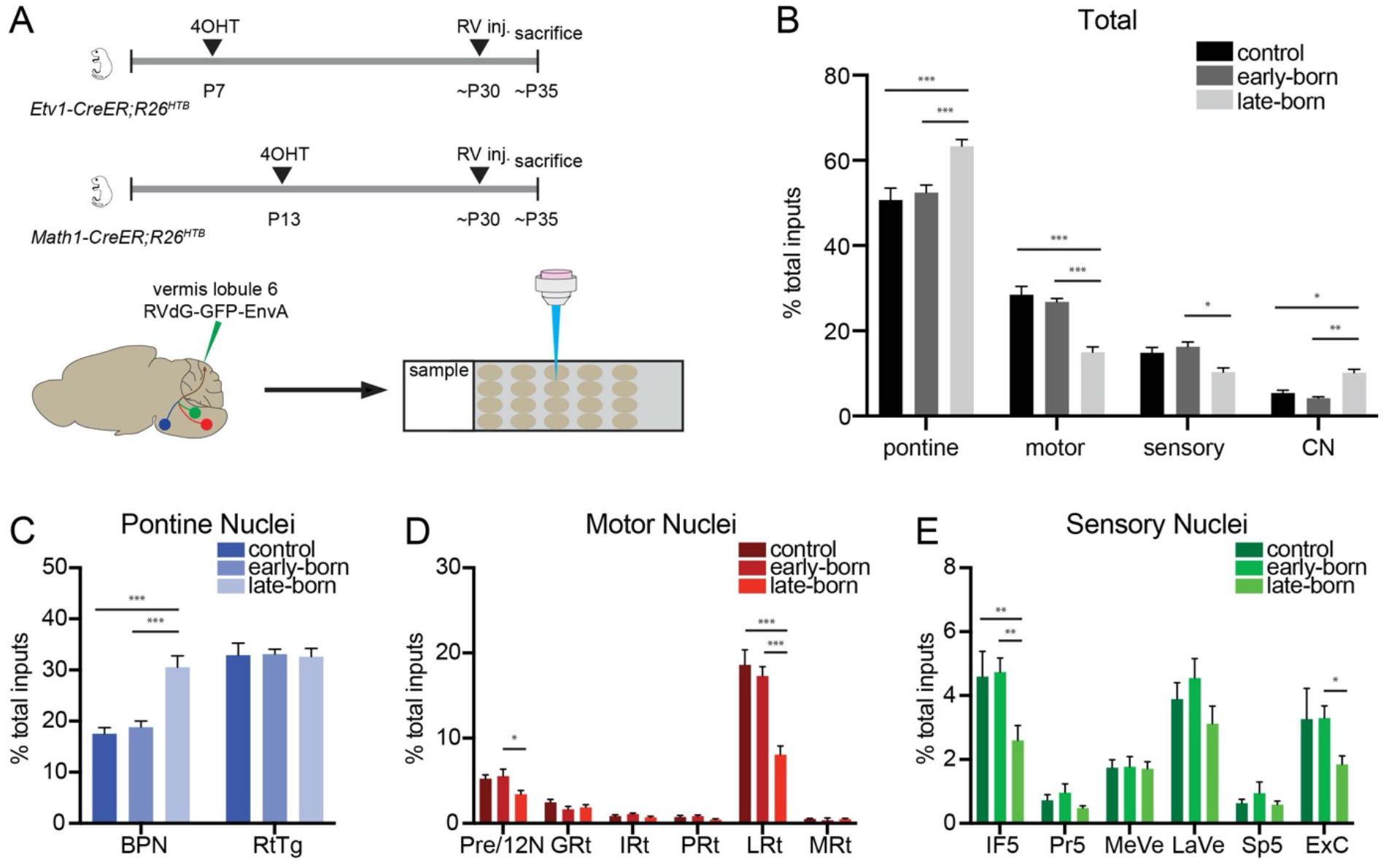
Rabies tracing of mossy fiber inputs to birth timing-defined vermis lobule 6 GrCs. (A) Schematic of experimental procedures; retrograde monosynaptic rabies tracing using RVdG-GFP-EnvA, rabies virus deleted for glycoprotein, expressing GFP, and pseudotyped with EnvA. (B) Total distribution of presynaptic inputs from super-categorized brainstem nuclei to control, early-born and late-born GrCs of vermis lobule 6. (C–E) Distribution of presynaptic inputs from brainstem pontine (C), motor (D), and sensory (E) nuclei to control, early-born and late-born GrCs of vermis lobule 6. N = 8 (control), 8 (early-born), 10 (late-born). Error bars, SEM. \**p* < 0.05, \*\**p* < 0.01, \*\*\**p* < 0.001 (ordinary two-way ANOVA with Tukey’s multiple comparisons test). See Figure 2 legend for anatomical abbreviations.

Quantification revealed that late-born GrCs receive more pontine and CN input than early-born GrCs, and less input from motor and sensory nuclei (Fig. 3B). Early-born GrCs received similar absolute and within-super-category proportions of input as control GrCs from all input sources, possibly due to more overlap between early-born and control GrCs (Fig. 1H, 3B–E, and S3). The increased pontine input received by late-born GrCs was due to an increase in BPN inputs (Fig. 3C). Late-born GrCs also received fewer Pre/12N, LRt, IF5, and ExC inputs than early-born GrCs (Fig. 3D and E). Overall, late-born GrCs received more of their within-pontine input from BPN than RtTg (Fig. S3A) and less of their within-motor nuclei input from GRt and LRt than early-born GrCs (Fig. S3B). These data suggest that late-born GrCs could represent a special population which receives a modified distribution of mossy fiber input or could reflect the greater specificity of our genetic access to late-born compared to early-born GrCs (Fig. 1H).

Taken together, these tracing results indicate that all precerebellar nuclei examined provide inputs to both early- and late-born GrCs, but with quantitative differences: sensory and motor nuclei provide more input to early-born GrCs, while basal pontine and cerebellar nuclei provide more to late-born GrCs.

### Both Early- and Late-born Granule Cells Encode Diverse Signals

To address whether GrCs born at different times differ in their response properties towards diverse stimuli, we designed a preparation allowing near-simultaneous 2-photon imaging of PF Ca^2+^ activity at two depths while mice perform an operant task (Fig. 4A–C), followed by presentation of a panel of appetitive, aversive, and neutral sensory stimuli (Fig. 4D; see Methods). To do so, we crossed *Etv1-CreER* mice to *Ai148* mice, which express high levels of GCaMP6f via tTA2/TRE-mediated transcriptional activation in a Cre-dependent manner (51). We injected *Etv1-CreER;Ai148* mice with 4OHT after P21 to drive GCaMP6f expression in a subset of PFs at all molecular layer depths without bias (Fig. 4A and Fig. S4A and B). We then trained water-restricted mice in an operant arm-reaching task with a randomly interspersed 20% of trials ending in reward (sucrose water) omission (10, 52). We imaged expert mice performing the operant task, followed by presentation of a stimulus panel containing visual and auditory stimuli, as well as a free reward and two aversive stimuli: orofacial air puff and tail shock; these five stimuli were repeatedly presented in a random order (Fig. 4D; see Methods). We imaged at two depths in the molecular layer, manually tuning the optical apparatus to identify the most superficial imaging field (~10 μm from the pial surface) and the deepest imaging field (~10 μm from the Purkinje cell layer) that we could access (Fig. 4C). This provided near-simultaneous access to Ca^2+^ activity of PFs arising from late- and early-born GrCs (Movie 1). We then registered PFs and correlated their responses with different portions of the task and with each stimulus (Fig. 4E and F). Superficial and deep PFs had similar Ca^2+^ transient rates (Fig. S4C). In total, we imaged 5 mice over 19 sessions with independent fields-of-view and thus recorded from 831 superficial molecular layer PFs and 511 deep molecular layer PFs from vermis lobule 6 and simplex.

**Fig. 4.**
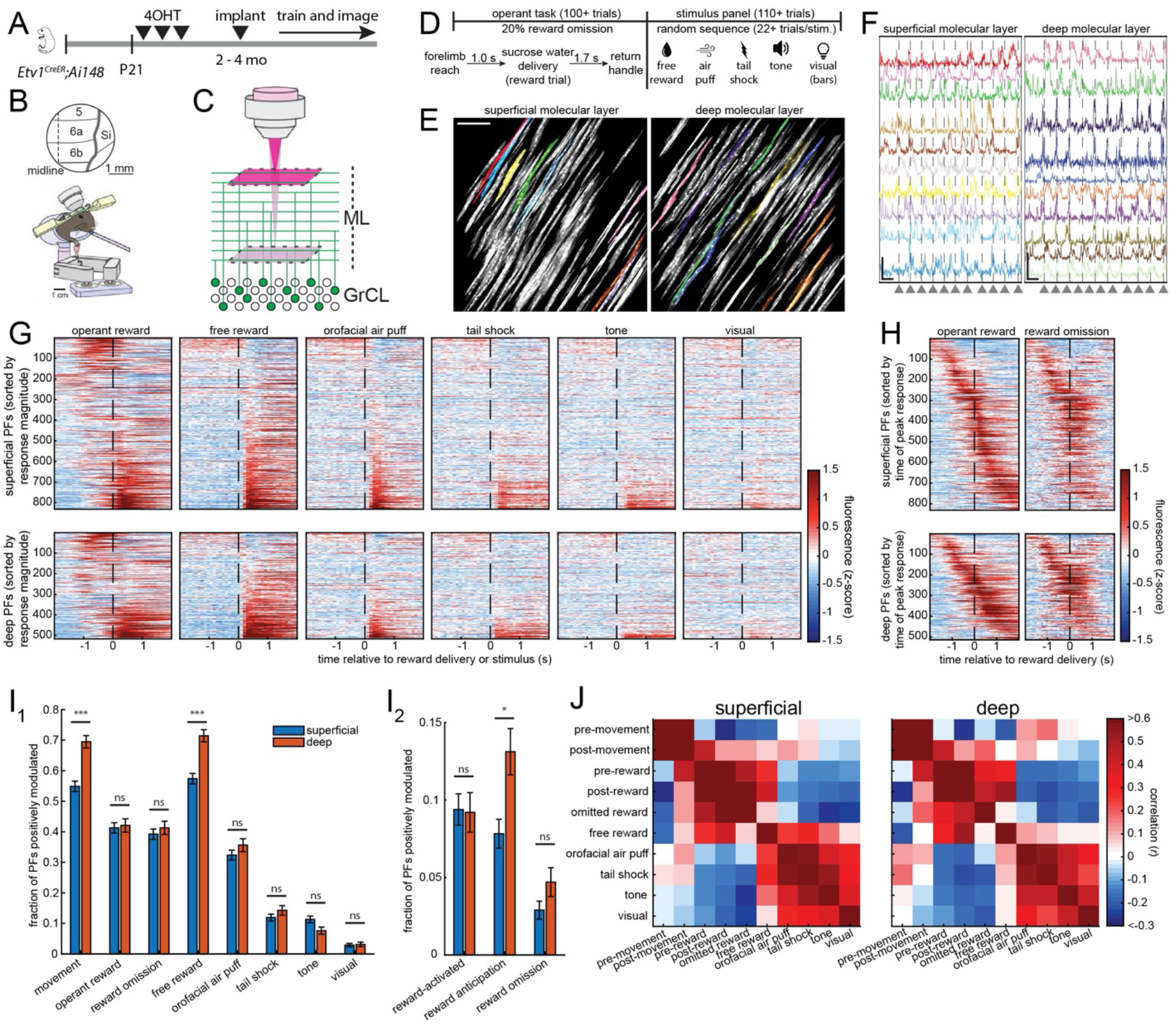
Simultaneous *in vivo* multi-depth two-photon imaging of deep and superficial parallel fibers (PFs) during an operant task and in response to a stimulus panel. (A) Genetic strategy for depth-unbiased GCaMP6 expression in PFs. (B) Window implantation. 5, vermis lobule 5; 6a/b, vermis lobule 6a/b; Si, simplex lobule. (C) Optical strategy for multi-depth two-photon near-simultaneous imaging of superficial and deep PFs, corresponding to late- and early-born GrCs, respectively. ML, molecular layer; GrCL, granule cell layer. (D) Operant task structure and stimulus panel with appetitive, aversive, auditory, and visual stimuli. (E) Example superficial and deep imaging fields-of-view with active PFs. Scale bar, 40 μm. (F) Example panels of PF Ca^2+^ responses from fields-of-view in panel D across several trials of the operant task. Gray arrowheads and vertical dashed lines denote reward onset. Vertical scale bars, 5 S.D. (standard deviation); horizontal scale bars, 5 seconds. (G, H) Heat plots of trial-averaged Ca^2+^ traces of individual PFs temporally aligned to reward delivery (G, H) or stimulus (G) onset and sorted by magnitude of response to operant reward or stimulus (G) or time of peak response during operant task trials (H) (831 superficial and 511 deep PFs from 19 sessions from 5 mice). (I) Fraction of PFs at both depths significantly positively modulated by each task variable or stimulus (H1; ***, *p* < 5 × 10^−6^ corrected; Wilcoxon rank-sum test with Holm-Bonferroni correction, n = 19 sessions), or by reward, reward anticipation, or reward omission (H2; *, *p* < 0.05 corrected; Wilcoxon rank-sum test with Holm-Bonferroni correction, n = 19 sessions). See Methods. (J) Linear regression analysis to explain each individual PF’s time-varying fluorescence as a weighted sum of behavioral events and stimuli (see Methods). Matrix shows the correlation coefficient between the weights assigned to each pairing of regressors across all PFs from either depth.

Overall, large fractions of both superficial and deep PFs exhibited responses to the operant task and free reward, smaller fractions responded to the aversive stimuli (orofacial air puff and tail shock), and even smaller fractions responded to the neutral auditory and visual stimuli (Fig. 4G). Both superficial and deep PFs encoded representations tiling the temporal scope of reward and reward omission trials in the operant task (Fig. 4H). Deep PFs, representing PFs of early-born GrCs, were positively modulated by movement and free reward in higher proportions than superficial PFs (Fig. 4I_1_ and Fig. S4F), but no such differences were found in PFs negatively modulated by the task variables and stimuli (Fig. S4D). We also recovered PFs activated by reward, reward anticipation, and reward omission (10); the deep molecular layer had a higher proportion of reward anticipation PFs than the superficial molecular layer (Fig. 4I_2_). Thus, while both superficial and deep PFs broadly represent task variables and stimuli, they do so in different proportions.

We also performed regression analysis using ten variables (pre-movement, post-movement, pre-operant reward, post-operant reward, reward omission, free reward, air puff, tail shock, tone, and visual stimuli) to uncover which variables best explain the dynamic activity levels of each PF (Fig. S5A and 5B). These analyses revealed that similar numbers of variables (regressors) contributed to activity changes in superficial and deep PFs (Fig. S5A), highlighting the potential for diverse, multimodal representations in PFs at each depth. This analysis also suggested that PFs tended to respond to clusters of different but potentially related task variables and stimuli, with two main clusters representing the task variables and stimuli (Fig. 4J). To estimate the dimensionality, or number of distinct modes of activity, in superficial and deep PF ensemble activity, we computed a principal components analysis on the complete imaging movies, and then tabulated the cumulative variance explained versus the number of principal components included in the reconstruction. Due to the difficulty of estimating dimensionality on a small number of cells, we alternatively computed the dimensionality on trial-averaged activity from all superficial or deep PFs in all imaging sessions. In both cases, a majority of the PF population activity variance was explained by ~5–8 principal dimensions (Fig. S5C).

### Evidence that Some Granule Cells Encode General Aversion

Acquisition of the Ca^2+^ activity traces of PFs in response to diverse stimuli enabled us to investigate their encoding properties further. We found that PFs activated by orofacial air puff were often also activated by tail shock (Fig. 5A and B). Comprehensive correlation analyses revealed a strong correlation between PFs responding to air puff and PFs responding to tail shock (Fig. 5B_1_, B_2_), representing the strongest correlation between the stimuli presented (Fig. 4J). By contrast, there was little correlation between free reward and air puff or tail shock (Fig. 5B_3_–B_6_). These trends were evident in PFs at both depths. As orofacial air puff and tail shock are unlikely to activate the same primary sensory responses, these data suggest that some GrCs could encode generalized aversive stimuli or punishment.

**Fig. 5.**
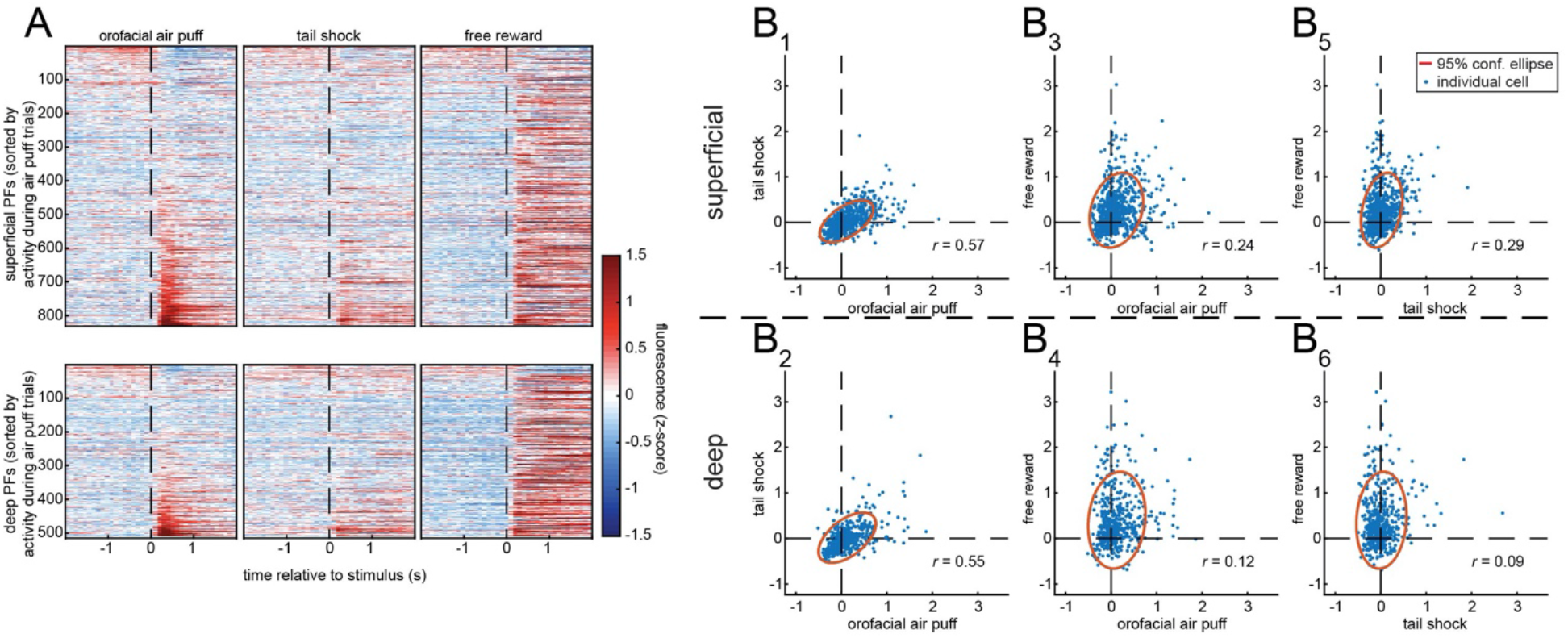
Evidence for GrC encoding of aversive stimuli. (A) Heat plot of trial-averaged Ca^2+^ traces of individual parallel fibers (PFs) temporally aligned to air puff and tail shock and sorted by magnitude of response to air puff (831 superficial and 511 deep PFs from 19 sessions from 5 mice). Following the analysis in Fig. 4J, for superficial (top) or deep PFs (bottom), correlations between the linear regression weight assigned to air puff versus tail shock (B_1_, B_2_), air puff versus free reward (B_3_, B_4_), and tail shock versus free reward (B_5_, B_6_). Each blue dot is a PF (831 superficial PFs, 511 deep PFs), and each red ellipse denotes 95% confidence. Pearson correlation coefficients (*r*) are displayed.

## Discussion

Birth timing is a determining factor for neuronal connectivity and function in a variety of systems from insects to mammals (25). In the mammalian neocortex, for example, birth timing dictates the cortical layers that glutamatergic excitatory neurons occupy (26, 27), which in turn predicts their connectivity patterns and functions (53). Birth timing also predicts the subtypes and cortical layers of GABAergic inhibitory neurons (54, 55) and the layer-specific axonal innervations of modulatory cholinergic neurons (29). In the cerebellar cortex, birth timing dictates the depth of the molecular layer to which the axons (PFs) of granule cells (GrCs) project (24). Even though PFs at different depths innervate the same Purkinje cells, whose dendrites span the entire molecular layer, there are at least two ways such spatially organized inputs could affect information processing. First, dendritic inputs at different distances from the soma can produce somatic membrane potential changes of different magnitudes and thus differentially affect action potential production (56, 57). Second, clustered synaptic inputs can summate nonlinearly to boost synaptic plasticity and transmission of synaptic potentials to the soma (56, 58–61). Thus, by mapping different mossy fiber populations to different depths of the Purkinje cell dendritic tree, GrC birth timing could modulate the impact of different inputs on cerebellar cortical output.

Our trans-synaptic tracing results indicate that early- and late-born GrCs receive inputs from the same collections of brainstem nuclei (Fig. 3). Our simultaneous *in vivo* imaging of deep and superficial PFs—corresponding to early- and late-born GrCs—indicate that they exhibit largely similar activity patterns during an operant task and in response to a battery of stimuli (Fig. 4). These data suggest that GrC input–output map is far from organized parallel processing, in which inputs from specific precerebellar nuclei are transmitted to specific depths of the molecular layer. Indeed, our imaging data largely agree with a recent report that did not detect differences in early- vs. late-born GrCs in their responses to sensorimotor stimuli (36). Together, these data are consistent with nonselective mossy fiber inputs onto GrCs born at different times, in line with the classic Marr-Albus framework (3, 4).

On the other hand, we identified quantitative differences between the distribution of mossy fiber inputs to early- vs. late-born GrCs, with brainstem sensory and motor nuclei contributing more inputs to early-born GrCs and pontine and cerebellar nuclei contributing more inputs to late-born GrCs. Likewise, our imaging identified small but significant differences in the fraction of GrCs positively modulated by movement, reward anticipation, and reward consumption. These data rule out a completely stochastic mossy fiber→GrC connectivity model. Future work is required to determine how differential mossy fiber inputs contribute to the differences in physiological response properties described here, and whether GrCs born at different times exhibit different activity patterns and functions in other behavioral paradigms.

The circuit architecture of the vertebrate cerebellar cortex has often been compared to that of the insect mushroom body (5, 62, 63). Both circuits feature dimensionality expansion: a smaller number of mossy fibers provides input to a much larger number of GrCs in the cerebellar cortex; a small number olfactory projection neurons (PNs) provides input to a much larger number of Kenyon cells (KCs, the intrinsic neurons of the mushroom body) for representation of different combinations of olfactory cues. Anatomical and physiological studies have provided evidence for random PN→KC connections (64–66), even though birth timing is an important determining factor for both PNs (67, 68) and KCs (69). However, other physiological studies and more recent serial electron microscopic reconstructions have uncovered non-random connectivity (63, 70–72). Specifically, a subset of axons of PNs that represent food odors converge onto individual KCs more frequently than expected from a random connectivity model; this property can be used to enhance discrimination of commonly encountered and ethologically relevant stimuli (72). Furthermore, three major KC classes born at different times from a common neuroblast receive biased input from different PN types (63). Altogether, these studies suggest that the mushroom body circuit utilizes random connectivity within local regions themselves nested within structured organization at a larger scale (63, 72). Whether mossy fiber→GrC connectivity follows similar rules awaits future investigation.

Our study also provides insights into GrC connectivity and coding unrelated to birth timing. Our description of mossy fiber inputs to vermis lobule 6 and simplex GrCs (Fig. 2) provide, to our knowledge, the first profiles of monosynaptic inputs to cerebellar GrCs. Notably, we found more pontine input to vermis lobule 6 than to simplex, a part of the cerebellar hemisphere (Fig. 2I). The vermis is classically considered a part of the spinocerebellar system, underscoring its reception of trigeminal and proximal somatosensory input; by contrast, the hemisphere is classically considered part of the cerebrocerebellar system, emphasizing its interconnections with the cerebral cortex (73–75). Our results suggest a different view with regards to inputs to vermis vs. hemisphere, and are in line with multi-synaptic retrograde tracing studies from the vermis of monkeys (48), rats (50), and mice (18, 49). Together, these studies and ours argue against a simple, dichotomous conception of mossy fiber input to vermis vs. hemisphere.

Recent physiological recording studies in behaving animals have highlighted the role of cerebellum in reward processing (10–12, 76–80). Specifically, individual GrCs can encode reward delivery, omission, or expectation (10). We found that a substantial number of GrCs are activated by orofacial air puff and tail shock (Fig. 5), two aversive stimuli that use distinct sensory pathways. This finding suggests that GrCs may also encode generalized aversiveness, complementing their representation of reward and expanding our understanding of the cerebellum’s role in cognitive processing (13).

## Materials and Methods

### Mice

All procedures followed animal care and biosafety guidelines approved by Stanford University’s Administrative Panel on Laboratory Animal Care and Administrative Panel of Biosafety in accordance with NIH guidelines. The day of birth was considered postnatal day 0 (P0). All mice were on CD1 and C57BL/6J mixed backgrounds. Transgenic lines *Etv1-CreER* (43), *Math1-CreER* (41), *Ai14* (44), *R26^HTB^* (45), and *Ai148* (51) were used where indicated. Mice used to demonstrate genetic strategies were perfused/sacrificed between P28 and P32. Mice used in anatomical experiments were injected with rabies virus between P28 and P33 and perfused 5 days later. Mice used in imaging experiments were implanted with imaging windows at between 10 and 14 weeks of age. Both males and females were used in anatomical experiments; females were used in imaging experiments. Mice were housed in plastic cages with disposable bedding on a 12-hour light-dark cycle with food and water available *ad libitum* until placed on water restriction.

### 4OHT production and injection

4-Hydroxytamoxifen (4OHT) was prepared by dissolving in Chen oil (81). *Math1-CreER;Ai14* and *Math1-CreER;ROSA26^HTB^* pups were injected intraperitoneally at P2 (50 mg/kg) or P13 (150 mg/kg). P13 *Etv1-CreER;Ai14* or *Etv1-CreER;ROSA26^HTB^* pups were injected intraperitoneally at P7 with 50–150 mg/kg 4OHT. *Etv1-CreER*;*Ai148* mice used in imaging experiments were injected intraperitoneally after P21 2–5 times with 150 mg/kg 4OHT.

### Immunostaining

Mice were deeply anesthetized using 2.5% Avertin and perfused transcardially, first with ~10 mL PBS then ~25 mL 4% paraformaldehyde (PFA) in PBS. The fixed brains were dissected out and postfixed overnight at 4°C in 4% PFA in PBS. Brains were then immersed in sucrose 24–48 hours, embedded in Optimum Cutting Temperature (OCT, Tissue Tek), and sectioned on a cryostat (Leica).

For genetic strategy proof-of-principle experiments, 40-μm sections of the cerebellar vermis were collected in PBS, washed twice in PBS for 10 minutes, incubated in 10% normal donkey serum (NDS) in PBST (PBS with 0.3% Triton X-100) for 2 hours at room temperature, and incubated in primary antibody (1:1,000 mouse anti-calbindin, Sigma; 1:500 rabbit anti-DsRed, Clontech; 1:1,000 guinea pig anti-vGluT1, Millipore) solution containing 5% NDS in PBST for 2 overnights at 4°C. Sections were then washed 3 times in PBST, each for 10 minutes; incubated in primary antibody solution containing 5% NDS in PBST for 2 hours at room temperature; washed in PBST for 10 minutes, incubated in PBST with DAPI (1:10,000 of 5 mg/mL, Sigma-Aldrich) for 30 minutes; then successively washed in PBST and PBS, each for 10 minutes. Sections were then mounted on Superfrost Plus slides, coverslipped with Fluoromount-G, and allowed to dry for at least 4 hours at room temperature before imaging.

For rabies tracing experiments, 60-μm sections encompassing the entire hindbrain (from AP −3.5 to −8.5) were collected onto Superfrost Plus slides in the order of sectioning. Slides were dried at room temperature overnight before further processing. Following overnight drying, slides were washed in PBST (PBS with 0.3% Triton X-100) for 10 minutes, incubated in PBST with DAPI (1:10,000 of 5 mg/mL, Sigma-Aldrich) for 30 minutes, then washed once in PBST and once in PBS, each for 10 minutes (all steps at room temperature). Slides were then dried and coverslipped with Fluoromount-G (Thermo Fisher Scientific). Slides were allowed to dry for at least 4 hours at room temperature before slide scanner imaging. Whole slides were then imaged with a 10x objective using a Leica Ariol slide scanner and a SL200 slide loader.

### Fluorescence Imaging

Images were taken on a Zeiss LSM 780 laser-scanning confocal microscope (Carl Zeiss). To quantify the portion of the molecular layer innervated by genetically labeled GrCs, fluorescence intensity measurements were taken on unprocessed images in Fiji (ImageJ) and data were processed using custom MATLAB scripts. A 400-pixel-wide segmented line was drawn from the border of the deep molecular layer to the pial surface (the border of the superficial molecular layer), as defined by counter stains with calbindin and vGluT1, and average intensity values along the line were measured using the Plot Profile command. The intensity traces were interpolated into 100 bins from deep to superficial using custom code (MATLAB), and the traces from individual sections were normalized to a maximum of 100. Numbers of sections were as follows: early-born strategy, 12 sections from 2 mice; late-born strategy, 12 sections from 2 mice; control strategy, 12 sections from 2 mice.

### Rabies Virus Production and Injection

Pseudotyped (EnvA-coated) G-deleted rabies virus (RV) encoding expression of eGFP were produced following a published protocol (82). Mice were anesthetized with 1.5–2.0% isoflurane and placed in a stereotaxic apparatus (Kopf Instruments). The following sets of coordinates (in mm) were used: 0.0 AP, 0.0 ML, –0.3 DV and 0.5 AP, 0.0 ML, –0.3 DV (relative to the posterior suture) for injections targeted to vermis lobule 6; and –2.0 AP, ±2.0 ML, –0.3 DV (relative to lambda) for injections targeted to bilateral simplex. For injections targeting control GrCs in vermis lobule 6 and simplex (Fig. 2), 100 nL RVdG-eGFP-EnvA was injected into each site. For injections targeting early- and late-born GrCs in vermis lobule 6 (Fig. 3), 300 nL RVdG-eGFP-EnvA was injected into each site. Following injection, mice were housed in a biosafety level 2 (BSL2) facility to allow for rabies transsynaptic spread and eGFP expression. All mice were injected between postnatal days 28 and 33 and sacrificed 5 days after RV injection.

### Quantification of Rabies Tracing Results

Quantification of brainstem subregions relied on boundaries outlined in a mouse brain atlas (83). Anatomically contiguous regions with similar functions and lacking clear-cut anatomical boundaries were combined. For example, the prepositus (Pre) and hypoglossal nuclei (12N) were combined due to their anatomical continuity, a lack of distinguishing anatomical markers in our histological preparation, and their co-involvement in motor coordination of facial muscles. The parvicellular reticular nucleus (PCRt) and its alpha part (PCRtA) were combined into PRt (parvicellular reticular nucleus); the intermediate reticular nucleus (IRt) and its alpha part (IRtA) were combined into IRt (intermediate reticular nucleus); the gigantocellular nucleus (Gi), including its alpha (GiA) and ventral (GiV) parts, and the lateral (LPGi) and dorsal (DPGi) paragigantocellular nuclei were combined into GRt (gigantocellular reticular nucleus); the dorsal (MdD) and ventral (MdV) parts of the medullary reticular nuclei were combined into MRt (medullary reticular nucleus). The magnocellular (MVeMC) and parvicellular (MVePC) parts of the vestibular nuclei and the vestibulocerebellar nucleus (VeCb) were combined into MeVe (medial vestibular nucleus); while the lateral (LVe), spinal (SpVe), and superior (SuVe) vestibular subnuclei were combined into LaVe (lateral vestibular nucleus). The dorsomedial (Pr5DL) and ventrolateral (Pr5VL) parts of the principal sensory trigeminal nucleus were combined into Pr5 (principal sensory trigeminal nucleus); and the oral (Sp5O), dorsomedial (DMSp5), interpolar (Sp5I), and caudal (Sp5C) parts of the spinal trigeminal nucleus were combined into Sp5 (spinal trigeminal nucleus). Regions contributing under 0.75% total input, including the spinal cord, nucleus of the facial nerve (7N), dorsal cochlear nucleus (DC), and inferior olive (IO) were not included in the final analysis due to the very low number of labeled cells in these regions.

### Tracing Statistics

Unpaired t tests with Bonferroni corrections and two-way ANOVAs with Tukey’s multiple comparisons tests were performed using Prism 9 (GraphPad).

### Window Implantation

Mice were anaesthetized using isoflurane (1.25–2.5% in 0.7–1.3 liter per minute of O2) during surgeries. Hair was removed from a small patch of skin, skin was cleaned, and an incision was made to remove the patch of skin. Connective tissue and muscle were then peeled back, and the skull was dried. A 3 mm diameter cranial window centered rostrocaudally over the post-lambda suture and 1.5 mm right of the midline was then exposed via drilling, positioning the window over cerebellar lobules VI and simplex. To seal the skull opening, a #0 3 mm diameter glass cover slip (Warner Instruments) was affixed to the bottom of a 3 mm outer diameter, 2.7 mm inner diameter stainless steel tube (McMaster) cut to 1 mm height. The glass/tube combination was stereotaxically inserted into the opening in the skull at an angle of 45° from the vertical axis and 25° from the AP axis. The window was then fixed in place and sealed with Metabond (Parkell). Next, a custom stainless steel head fixation plate was fixed to the skull with Metabond and dental cement (Coltene Whaledent). The 1.2 mm thickness fixation plate had a 5 mm opening to accommodate the stainless steel tube protruding from the window, and two lateral extensions to permit fixing the plate to stainless steel holding bars during imaging and behavior.

### Operant Task

Briefly, mice were trained as described previously (10). During water restriction, mice were monitored daily for signs of distress, coat quality, eye closing, hunching and lethargy to ensure adequate water intake. Following two days of water restriction, mice were trained for 10–14 days for about 20–60 min daily, depending on performance and satiety. In both tasks, we recorded licking at 200 Hz using a capacitive sensor coupled to the metal water port which delivered approximately ~6 μl 4% sucrose water reward near the animal’s mouth. During training and in all experiments, mice were head-fixed, with their bodies from the torso down in a custom printed optically transparent plastic tube. Mice learned to voluntarily initiate pushing the handle of a manipulandum. The robot constrained movement of the manipulandum to the forward axis. Handle position was controlled and monitored by two motors and encoders (Maxon B7A1F24007CF, containing DCX22S EB KL 24V motor with ENX 16 RIO 65536IMP encoder), and robotic control relied on nested feedback loops in FPGA (10 kHz) and a real-time operating system computer (1 kHz), both in a National Instruments cRIO chassis, as well as a Windows PC (200 Hz). The controllers were programmed in LabVIEW and permitted precise robotic positioning and application of forces to the handle with a 1 kHz bandwidth to restrict motion as needed (52). The device recorded the handle position with a 200 Hz sampling rate and encoder resolution of 0.003 mm and permitted linear movements of maximum length 8 mm, after which the trial terminated. Following a delay (1 s), a solenoid delivered the water reward. Following another delay (1.7 s) the handle began to return to the home position. This process took 2 s to complete, after which the mouse could initiate the next movement at any time. During reward omission trials, reward was withheld on a randomly interspersed 20% of trials, but never on two consecutive trials.

### Imaging and Optics

All Ca^2+^ imaging was performed using a 40× 0.8 NA objective (LUMPlanFLN-W, Olympus) and a custom two-photon microscope with an articulating objective arm. 920 nm laser excitation was delivered to the sample from a Ti:sapphire laser (MaiTai, Spectra Physics) at powers of around 40/50 mW at the superficial and deep imaging depths, respectively. ScanImage software (Vidrio Technologies) was used to control all image acquisition hardware. In order to image the superficial and deep molecular layer together, suitable depths for each were first identified. A z-piezo (P-725.4CD, Physik Instrumente) was used to lower the focal plane just below the pial surface until a full-field plane of the molecular layer was visualizable; this plane was recorded as the superficial depth. A full-field plane of the molecular layer just above the Purkinje cell layer (40 – 100 μm below the superficial field-of-view) was then focused to and recorded as the deep depth. During imaging, ScanImage interleaved frame acquisitions at each depth by moving the z-piezo between frames, yielding a volume acquisition rate of 9.75 Hz. At each depth, resonant galvanometers scanned a 240 × 240 μm field-of-view. To ensure alignment of the articulating objective to the glass window on the brain, a back-reflection procedure was performed. A low power visible red laser (CPS180, ThorLabs) co-aligned to the infrared beam onto the glass window was projected into the cerebellar window; the red back-reflection was then viewed via an iris placed on the objective port. The mouse and objective angles were positioned to center the back-reflection into the iris aperture. During image acquisition, slow axial drifts were compensated in real time by frequently comparing the acquired images to initial images and correcting using an objective *z*-piezo (P-725.4CD, Physik Instrumente). To align imaging data to behavioral data, the behavioral computer acquired the microscope’s frame clock signal simultaneously with each mouse’s behavioral data.

### Stimulus Panel

A stimulus panel including delivery of free reward, orofacial air puff, tail shock, sliding tone and visual stimulus was designed. A total of 110 to 120 stimuli were presented over 15.4 minutes, with equal time intervals between presentation of successive stimuli. Free rewards consisted of a water reward of the same size as those of the operant task. Orofacial air puffs aimed at the mouse’s left whisker pad and eye were triggered through an external custom-built stimulus delivery system; air puffs were tested before imaging and always elicited eyeblink. Tail shocks were administered via an electrical stimulator (model #A320D, World Precision Instruments); electrode gel (Spectra 360, Parker Laboratories) was placed on the tail and the positive and negative leads (18-ga copper wire) were securely taped onto the middle portion of the tail 1 cm apart, and a 5V pulse to the stimulator triggered delivery of an electrical current (5 mA, 2 s). The auditory tone stimulus consisted of a wav file encoding a 2-second-long tone rising from 1 kHz to 20 kHz and then back down to 1 kHz and was delivered via speakers from 1 m away from the mouse. The visual stimulus consisted of a 1 s duration moving vertical bar of blue light on a custom-built LED grid (Sparkfun WS2812B).

### Image Processing

Files containing both superficial and deep images were first split by depth. Normcorre image registration software corrected rigid and piecewise-nonrigid lateral brain motion (84). Downsampled videos were manually inspected, and imaging sessions with significant brain motion/depth shift were discarded. Individual active PFs were identified in our imaging videos using automated cell sorting based on principal and independent component analyses (PCA/ICA). PFs corresponded to a weighted sum of pixels forming a spatial filter. Automated and manual segmentation and thresholding were used to truncate these filters down to individual PFs by eliminating spurious, disconnected components. Each PF’s time-varying fluorescence trace was extracted by applying the spatial filter to the processed videos. Slow drift was removed from each trace by subtracting a tenth-percentile-filtered (15 s sliding window) version of the signal. Finally, each PF’s fluorescence trace was *z*-scored to correct for differences in brightness between PFs; all fluorescence values were then reported in SD units. Data was aligned to the time of reward or stimulus delivery. For reward omission trials, data was aligned to the time at which reward would have been delivered following movement termination. The aligned fluorescence response of each PF was averaged across trials to produce the triggered averages shown.

### Imaging Statistics

MATLAB (Mathworks) was used for all statistical tests. All comparisons of unpaired means of two groups used Wilcoxon rank sum tests. All comparisons of paired means of two groups used Wilcoxon signed rank tests. *p* values were adjusted using the Holm-Bonferroni multiple comparisons correction.

### Modulation Analysis

Time windows for modulation analysis were defined with reward (or, for reward omission trials, when reward would have been delivered) or stimulus delivery timepoints serving as timepoint 0.0 s. To determine whether a PF was significantly modulated by a task variable/stimulus, we compared fluorescence before and after stimulus/variable. For each operant trial, we computed fluorescence for each cell averaged over the two time windows [0.0, 0.5] s and [-0.5, 0.0] s, while for each stimulus delivery we used the time windows [-1.0, 0.0] s and [0, 1.0] s. We then compared fluorescence in the two time windows. Significance for each cell was determined using a cutoff of *p* = 0.05.

For the operant task, we defined reward-activated PFs as those whose fluorescence was significantly greater after than before reward, and significantly greater after reward than after reward omission (*p* < 0.05 for both comparisons). We defined reward anticipation PFs as those whose fluorescence was significantly greater after reward omission than after reward, and significantly greater just prior to reward than prior to movement (*p* < 0.05 for both comparisons). We defined reward omission PFs as those whose fluorescence was significantly greater after reward omissions than after reward (*p* < 0.05), excluding those previously defined as reward anticipation PFs. In analysis of these three PF response types, pre and post time windows were taken to be [0.0, 0.5] s and [−0.5, 0.0] s.

### Regression Analysis

To determine which task variables/stimuli predict PF activity, we used linear regression analysis to reproduce the time-varying fluorescence of each individual cell as a weighted sum of boxcar “indicator” functions corresponding to each behavioral variable of interest. The operant regressors corresponding to pre-movement, post-movement, pre-reward, and post-reward used boxcars of width 0.5 s just prior or subsequent to either movement or reward. The regressor corresponding to operant reward omission used a boxcar in the later period, [0.5, 1.5] s with respect to reward omission. The regressors corresponding to the stimuli—free reward, air puff, tail shock, tone, and visual—used boxcars at [0, 1] s with respect to stimulus onset. These coefficients were defined as significant if *p* < 0.01.

### Principal Components Analysis

To quantify the number of dimensions in the PF imaging data, we computed principal components analysis (PCA). For a single-session and single-trial analysis, we took the entire T×N movie matrix (T timepoints, N parallel fibers) and computed PCA. For an across-sessions, trial-averaged analysis, for each of the 6 contexts (operant reward, free reward, puff, shock, tone, visual) we computed the trial-averaged response (consisting of T_trail_ timepoints) for all PFs, then concatenated all trial types into a single 6T_trail_×N_total_ matrix (where N_total_ is the total number of PFs recorded from all sessions and mice). For either single-trial or trial-averaged, we computed PCA on the data matrix and tabulated the cumulative variance explained as a rising function of the number of principal components included.

### Event detection

Ca^2+^ transients were defined as fluorescence local maxima (highest peak within 0.5 s) of minimum magnitude 1.5 z-scores.

## Data availability

Reagents and code are available upon reasonable request to the corresponding author.

## Acknowledgements

We thank the Luo lab, especially H. Li, J. Lui, and Y. Takeo, for feedback on the project and comments on the manuscript; M. Molacavage for administrative assistance; C. Manalac for assistance with genotyping; and A. Joyner for the *Math1-CreER* mice and M. Goulding for *R26^HTB^* mice. S.A.S. was supported by a National Science Foundation Graduate Research Fellowship and a Regina Casper Stanford Graduate Fellowship. M.J.W. was supported by a Career Award at the Scientific Interface from the Burroughs Wellcome Fund. N.P-D. was supported by the Biology Summer Undergraduate Research Program and an Undergraduate Advising and Research Major Grant. K.T.B. was supported by the Brain and Behavior Research Foundation (NARSAD 26845). L.L. is an investigator of the Howard Hughes Medical Institute. This work was supported by National Institutes of Health grants K99-DA041445 (to K.T.B.) and R01-NS080835 (to L.L.).

## Author Contributions

S.A.S. conceived the project, designed and performed all experiments and analyzed data. M.J.W. contributed substantially to all aspects of the 2-photon imaging experiments, including design, implementation, and data analysis. N.P-D. assisted with pilot tracing experiments. J.R., K.T.B. and S.M.G. provided viral reagents. T.H.K. and M.J.S. provided critical equipment and assistance with the 2-photon imaging experiments. S.A.S., M.J.W., and L.L. wrote the paper. L.L. supervised the project.

---

**Fig. S1.**
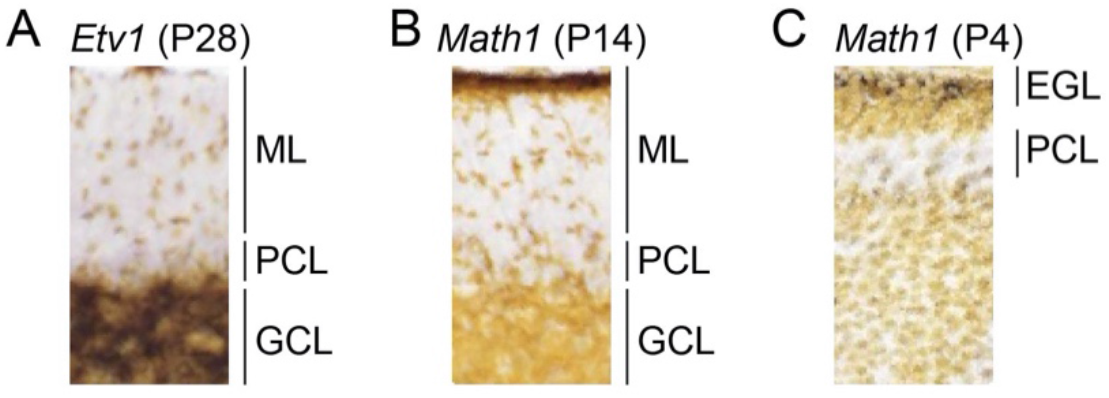
Expression patterns of transcription factors *Etv1* and *Math1*. (A–C) *In situ* hybridization of *Etv1* at postnatal day 28 (P28, A), *Math1* at P14 (B), and *Math1* at P4 (C) in cerebellar cortex vermis lobule 6. Data are from Allen Brain Atlas (40). ML, molecular layer; PCL, Purkinje cell layer; GCL, granule cell layer; EGL, external germinal layer.

**Fig. S2.**
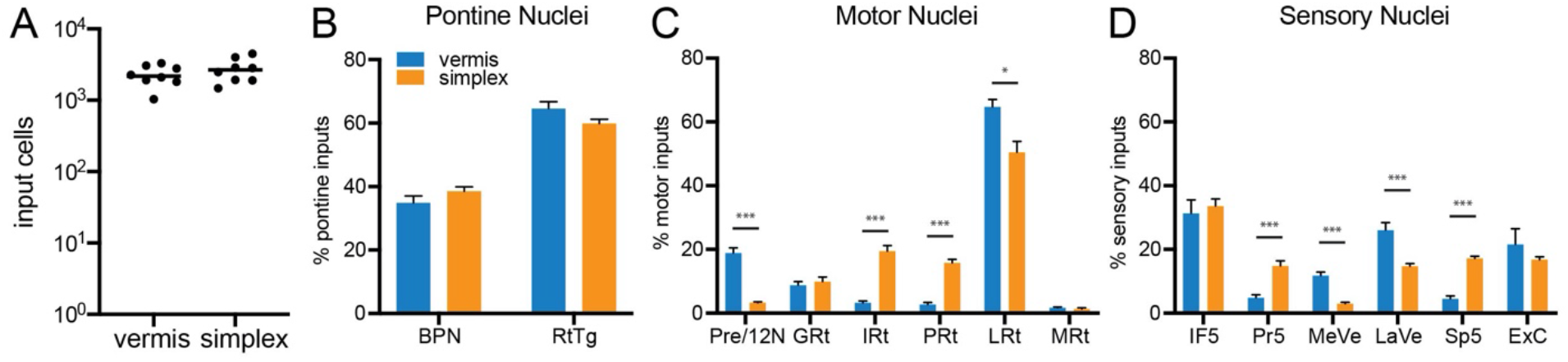
Additional analysis of mossy fiber inputs to granule cells (GrCs) of vermis lobule 6 and simplex. (A) Number of input cells to vermis lobule 6 and simplex control GrCs not defined by birth timing. (B–D) Proportion of inputs to vermis lobule 6 and simplex control GrCs contributed by each nucleus to total inputs from brainstem pontine (B), motor (C), and sensory (D) regions. N = 8 (vermis), 8 (simplex). Error bars, SEM. \**p* < 0.05, \*\**p* < 0.01, \*\*\**p* < 0.001 (multiple unpaired t tests with Bonferroni correction). See Figure 2 legend for anatomical abbreviations.

**Fig. S3.**
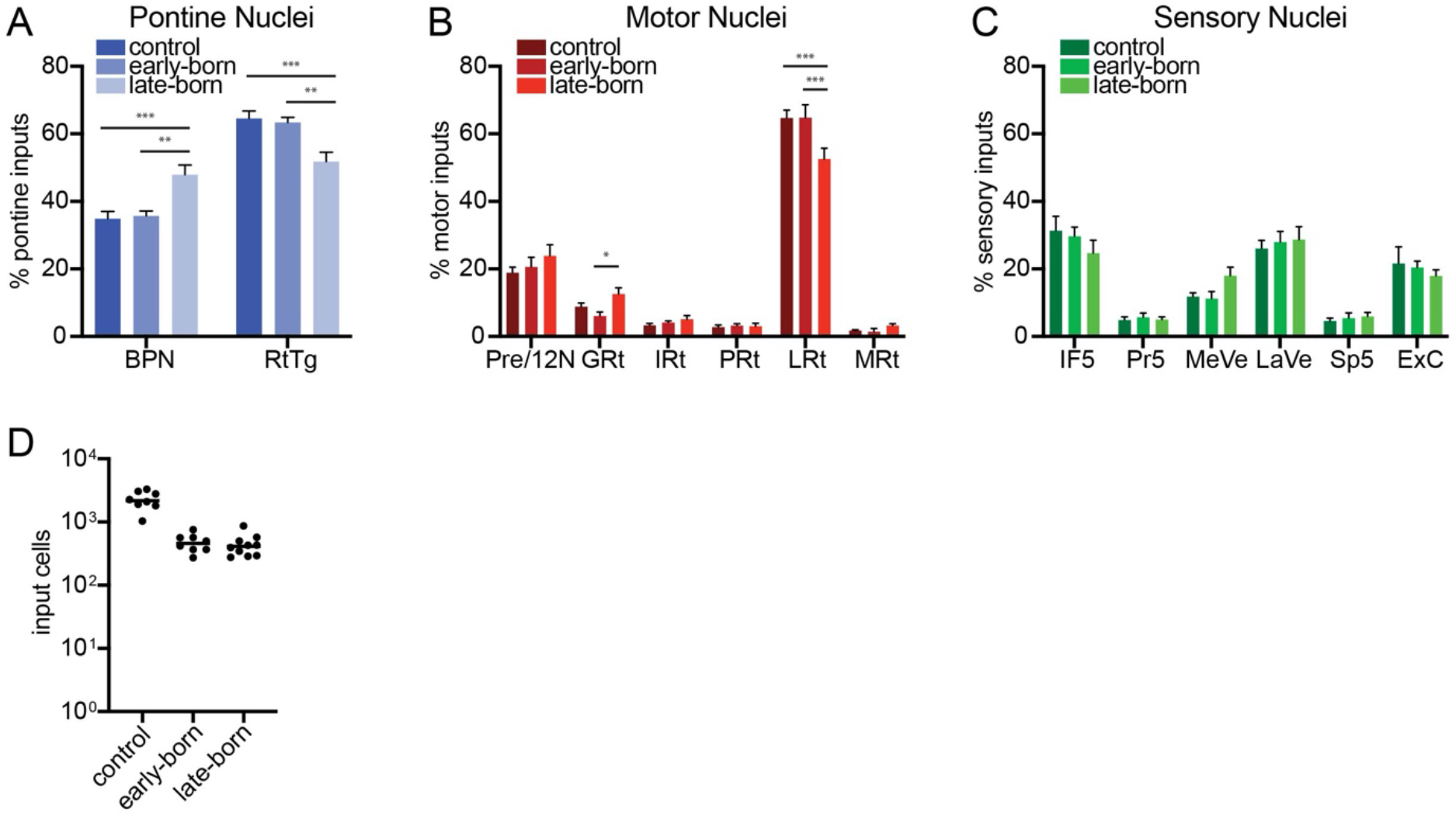
Additional analysis of mossy fiber inputs to birth timing-defined vermis lobule 6 GrCs. (A–C) Proportion of inputs to vermis lobule 6 control, early-born, and late-born GrCs contributed by each nucleus to total inputs from brainstem pontine (B), motor (C), and sensory (D) regions. N = 8 (control), 8 (early-born), 10 (late-born). Error bars, SEM. \**p* < 0.05, \*\**p* < 0.01, \*\*\**p* < 0.001 (ordinary two-way ANOVA with Tukey’s multiple comparisons test). (D) Number of input cells to vermis lobule 6 control, early-born, and late-born GrCs. See Figure 2 legend for anatomical abbreviations.

**Fig. S4.**
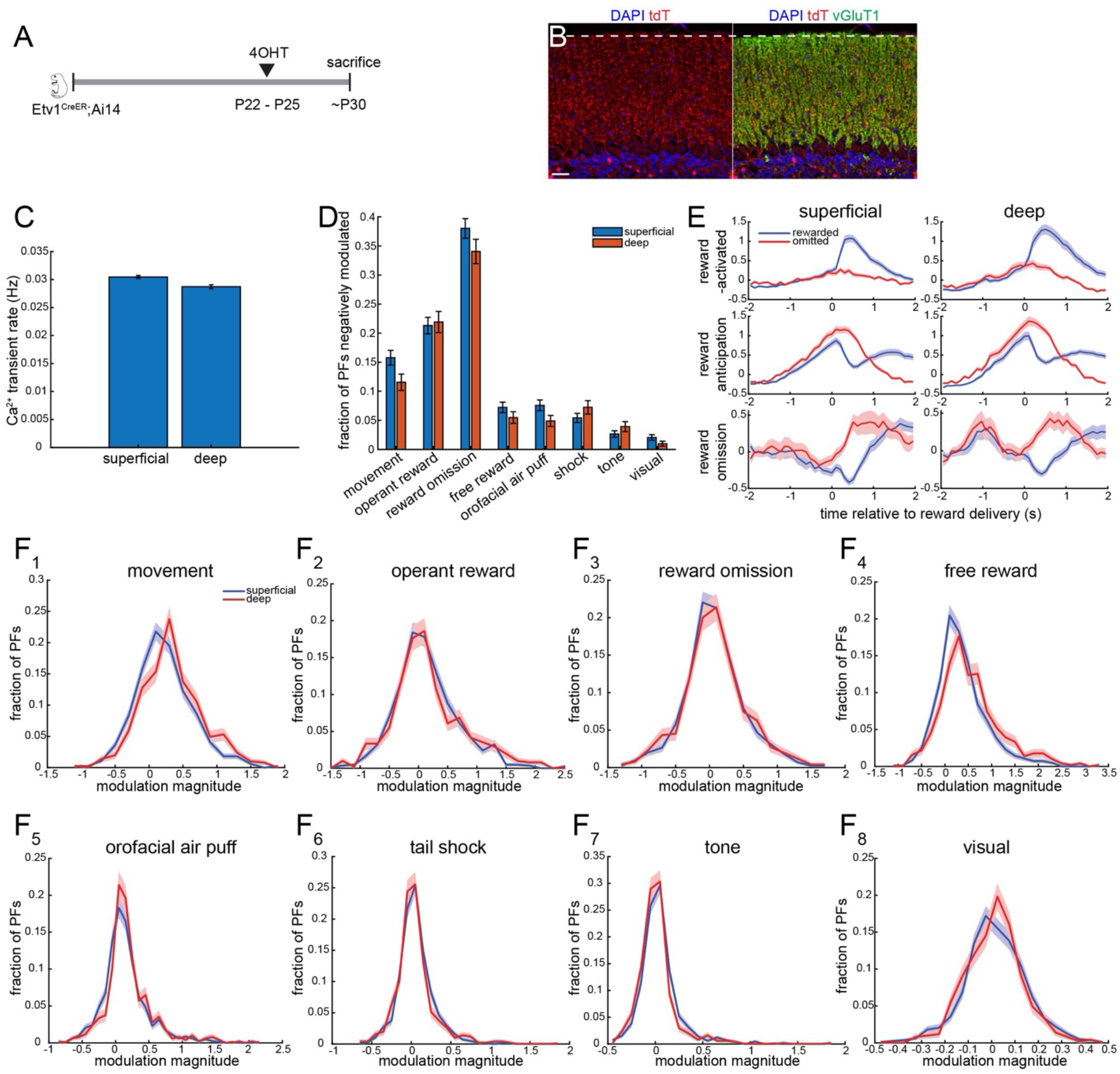
Additional analysis of multi-depth two-photon imaging data, Part I. (A) Genetic strategy for expressing GCaMP6f sparsely in GrCs not defined by birth timing, producing PFs that span all depths. (B) Fluorescence images showing PF innervation of the cerebellar molecular layer in a sparse and birth-timing agnostic manner. Image was taken from vermis lobule 6a. Scale bar, 20 μm. (C) Ca^2+^ transient rates of PFs at both depths. (D) Fraction of PFs at both depths significantly negatively modulated by each task variable or stimulus. Error bars, SEM; all pairs ns, *p* > 0.05 (Wilcoxon rank-sum test with Holm-Bonferroni correction, n = 19 sessions). (E) Averaged traces of PFs at both depths identified as reward-activated, reward anticipation, or reward omission PFs. (F) Distribution of modulation magnitudes of PFs at both depths by each task variable or stimulus.

**Fig. S5.**
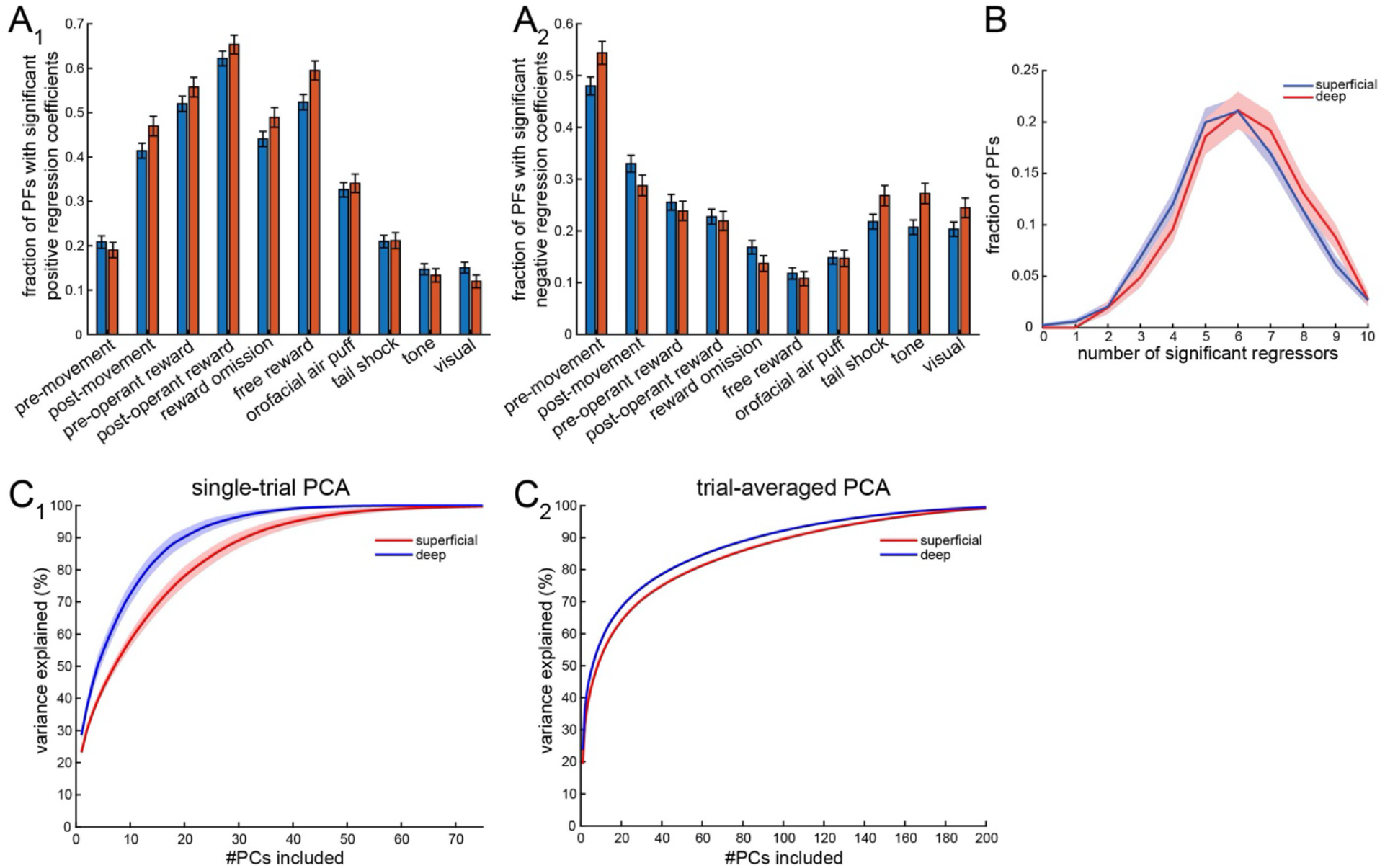
Additional analysis of multi-depth two-photon imaging data, Part II. (A) Using the regression analysis described in Fig. 4J, fraction of PFs at both depths with significant positive (A_1_) or negative (A_2_) regressors for each task variable or stimulus. Error bars, SEM; all pairs ns, *p* > 0.05 (Wilcoxon rank-sum test with Holm-Bonferroni correction, n = 19 sessions). (B) Distribution of number of significant regressors of PFs at both depths. Error bars, SEM (n = 19 sessions). (C) Principal components analysis (PCA) of superficial and deep PF ensemble activity based on dimensionality computed from single-trial (C_1_; mean ± SEM, n = 19 sessions) or trial-averaged (C_2_) activity of all superficial or deep PFs in all imaging sessions (see Methods). In both cases, about 5–8 dimensions are needed to explain half of the ensemble variance.

**Movie 1**: Multi-depth, near-simultaneous Ca^2+^ imaging of parallel fibers in the superficial (left) and deep (right) molecular layer during operant task. Movie is 4× temporally downsampled and is thus played at 4x speed. Original frame rate is 9.75 Hz.

